# Combined effect of biological and physical stress on artificial production of agarwood oleoresin in *Aquilaria malaccensis*

**DOI:** 10.1101/2020.05.19.103671

**Authors:** Hemraj Chhipa, Nutan Kaushik

## Abstract

Agarwood is the most expensive wood of the world and highly demanded wood in perfumery industry and ritual ceremonies of various religions. Agarwood is the infectious wood part of *Aquilaria* tree. Naturally, production of agarwood in *Aquilaria* takes 10-20 years of time and it can develop only in 1-2% of *Aquilaria* trees. Different types of biological, chemical and physical methods have been developed for artificial production of agarwood to fulfil the rising demand of the market. In the current article, we tried to explore combined effect of physical and biological stress in the form of stick method to improve agarwood production in *Aquilaria malaccensis* and compared it with well-known artificial fungal infection syringe method. Total 21 fungal strains were applied alone (syringe method) and with bamboo sticks (stick method). We found maximum infection occurred in stick method by fungi *Penicillium polonicum* AQGGR1.1 with 10 cm infection length. Artificial induction of marker compounds of agarwood, benzyl acetone and anisyl acetone were measured mostly in stick method, induced by 71.4% fungal strains grown on bamboo sticks, while alone only 42.9% fungi can induced in syringe method. *Penicillium aethiopicum* AQGGR1.2 found highly agarwood oleoresin inducing fungus in stick method and shown high potential agent in stick method for artificial production of agarwood.

## 1. Introduction

Agarwood is the infectious dark coloured wood part of tree *Aquilaria malaccensis* which contain various type of volatile compounds in agarwood oleoresin. Agarwood oleoresin composed of different types of Sesquiterpene and Chromone derivatives, and their composition determines the quality or grade of agarwood in the market. Presence of Sesquiterpene and Chromone mainly responsible for particular aroma and pleasant odor of agarwood (Takemoto et al. 2008; Wetwitayaklung et al. 2009). In Sesquiterpene, Agarofuran, Agarospiranes, Guaianes, Eudesmanes, Eremophilanes, Prezizaanes; while Chrome derivatives 2-(2-Phenylethyl)-4H-chromen 4-one, Di-epoxy tetra hydro chromones aromatics compounds and Triterpenes has been explored in different species of *Aquilaria* (Chen et al. 2012; Kalra and Kaushik 2017; Ueda et al. 2006; Ishihara et al.1991 and 1992). The value of agarwood has been estimated in the range of 9700 to 32000 USD per Kg, which is depends on quality grade of agarwood (Sen et al. 2015). Naturally, oleoresin production in Aquilaria tree is the result of the wound deterrence caused by microbial or insect infection or physical injury, which takes several years in its production (Blanchette & Van Beek 2009; Pojanagaroon and Kaewrak 2005; Gerard 2007). It is a kind of plant defence response against infection or injury and plant secretes oleoresins to prevent them, caused by physical and biological agents. Economic importance in the pharmaceutical and perfumery industry of quality grade of oleoresin makes the Aquilaria wood high priced commodity (Nor Azah et al. 2008). The market of agarwood is increasing rapidly and expected around 6-8 billion USD (Sen et al. 2015). Different artificial infection methods have been developed to increase the production of oleoresin (Mohamed et al. 2010; Zhang et al. 2012; Chhipa et al 2017). Nature of artificial infection method affects the diversity of compounds in oleoresin and host pathogen interaction also responsible for metabolites composition of the host (Nema 1989; Prasad et al. 1988; Khatri et al. 1985). Wounding by physical methods generate inferior quality agarwood which do not satisfy market demand (Persoon 2007).

Recently, Liu et al. (2013) developed whole tree agarwood inducing technique (Agar wit) which improved agarwood yield 4 to 28 times more. Further, in biological methods, fungi also explored in artificially production of agarwood (Tamuli et al. 2005; Tian et al. 2013; Zhang et al. 2012; Chhipa and Kaushik 2017). Hitherto, there are gaps in role of fungal stains in chemical composition modification of artificially developed agarwood which needs to be explored for best quality agarwood production by artificial infection. In our previous study we have reported use of syringe method for induction of artificial infection (Chhipa and Kaushik, 2017). In the present study, we tried to develop combined biological (fungi) and physical (stick) injury to increase stress on plant for more oleoresin production. We tried to identify the responsible fungi and effect of combined stress for higher production of marker compounds in agarwood oleoresin.

## 2. Materials and methods

Four to seven years old *Aquilaria malaccensis* trees in Dergaon (26.7°N 93.97°E), Assam and Jairampur (27°21′4″N 96°0′57″E), Arunachal Pradesh, India were used for artificial infection experiments. Identification of *Aquilaria malaccensis* species was confirmed by Dr. LR Bhuyan, Botanist, State Forest Research Institute Itanagar, Arunachal Pradesh before artificial infection. The fungal strains used in stick and syringe method were isolated from *Aquilaria malaccensis* and circumventing soil of *Aquilaria malaccensis* trees and identified by amplification of their ITS regions and similarity analysis of obtained ITS sequence with NCBI database according to Chhipa and Kaushik (2017). Fungal strains were maintained by routine sub-culturing on PDA medium for further experiments.

### 2.1 Artificial infection methods

#### 2.1.1 Stick method

In stick method, bamboo sticks (10 cm × 0.5 cm L × W) were obtained from TERI Gram, Haryana, India. The sterilized sticks were added to 2 days old grown fungal culture on potato dextrose broth in 250 ml of culture jars and incubated for 7 days at 25 ± 2°C. After growth of fungi on sticks, they were transferred to sterile falcon tube for experiment purpose. The holes in *Aquilaria* tree were drilled up to 8 cm in depth by sterilized drilling machine and holes were filled with fungal grown stick per hole. The sticks were inserted up to 5 cm. Each strain contain sticks were inserted in replicates and uninfected bamboo sticks were used as control. The infected wood around sticks was collected after 3 months of incubation and oleoresin compounds were extracted by refluxing method and analyzed by Gas Chromatography-Mass Spectroscopy (7890A/5975C, Agilent) according Chhipa and Kaushik (2017). Figure1 shown various steps involved in the procedure.

**Figure. 1:**
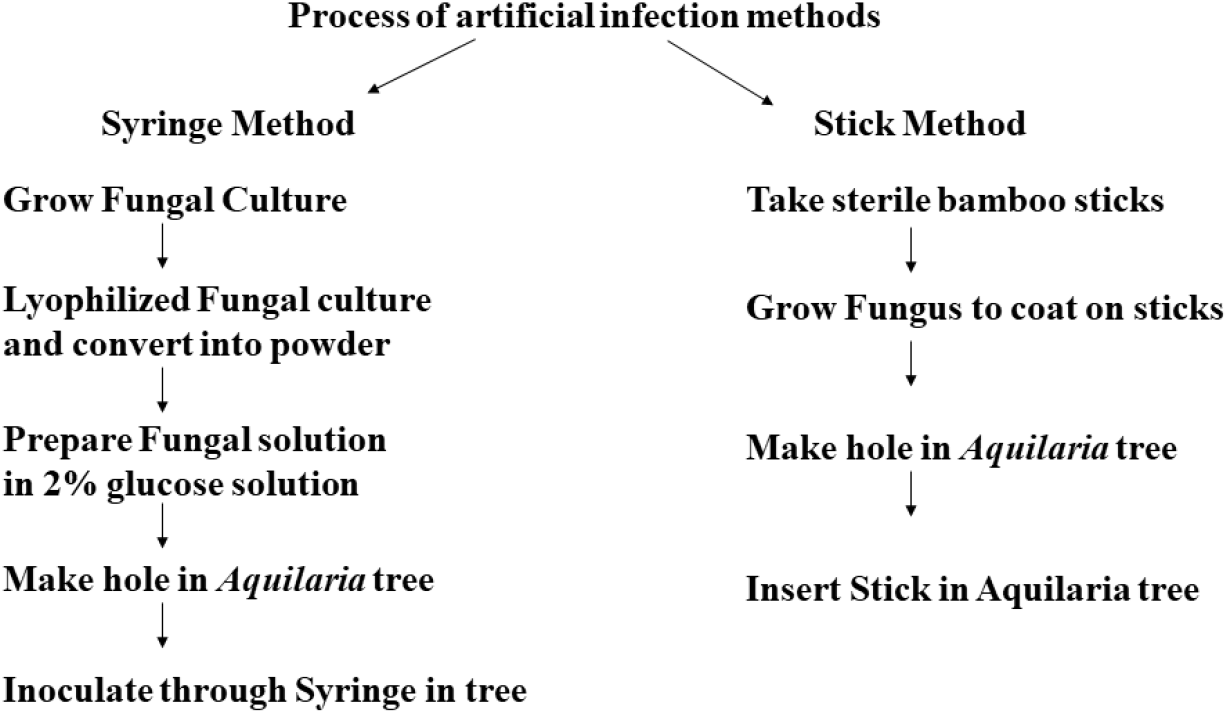
Process of artificial infection methods

#### 2.1.2 Syringe method

Artificial inoculation via syringe method was done according to Chhipa and Kaushik (2017). Briefly, seven days grown fungal biomass was lyophilized and converted into powder. The syringe inoculum was prepared in 2% glucose solution at 1 mg/ml concentration at the site of experiment. The fungal solution was injected in holes through syringe, which were drilled by sterilized drilling machine in zig-zag manner (Figure 1). First wound inflicted at 20 cm above the ground and next wound drilled at 10 cm interval. The hole was drilled up to 1.5 cm in depth and 1 ml of fungal inoculum was injected into 1.5 cm hole by sterile syringe and hole was sealed with parafilm. 2% glucose solution was used as control and each fungal strains were injected in replicates. Similarly, infected wood was collected and analyzed by Gas Chromatography as described in stick method.

## 3. Results

### 3.1. Effect on infection length

The effect of fungi on wound length was measured after 3 months of infection, by scrapping the bark of the tree around infected holes. It was observed that wood color turned from white to yellow or dark brown. Different fungal strains showed variation in wound formation in each method. In stick method maximum infection length was measured up to 10.00 cm caused by *Penicillium polonicum* AQGGR1.1 but in syringe method it was infected only 2.6 cm in length. Similarly, in syringe method maximum wound infection was found 8.5 cm caused by *Aspergillus flavus* strain AQGSS10 and it could infect only 2.83 cm in stick method (Figure 2). Different strains of fungal species also showed variation in wound infection length. Various *Aspergillus* species used in this study have shown wound formation in the range of 2 to 8.5 cm of infection length, while various *Penicillium* species infected *Aquilaria* trees in the range of 2 to 10 cm. The infection length in all treated trees were monitored up to 6 months. In stick method infection was increased up to 133.7% while in syringe method, only 49.7% was increased after 3 months of development of agarwood oleoresin initiation (Table S1).

**Figure. 2:**
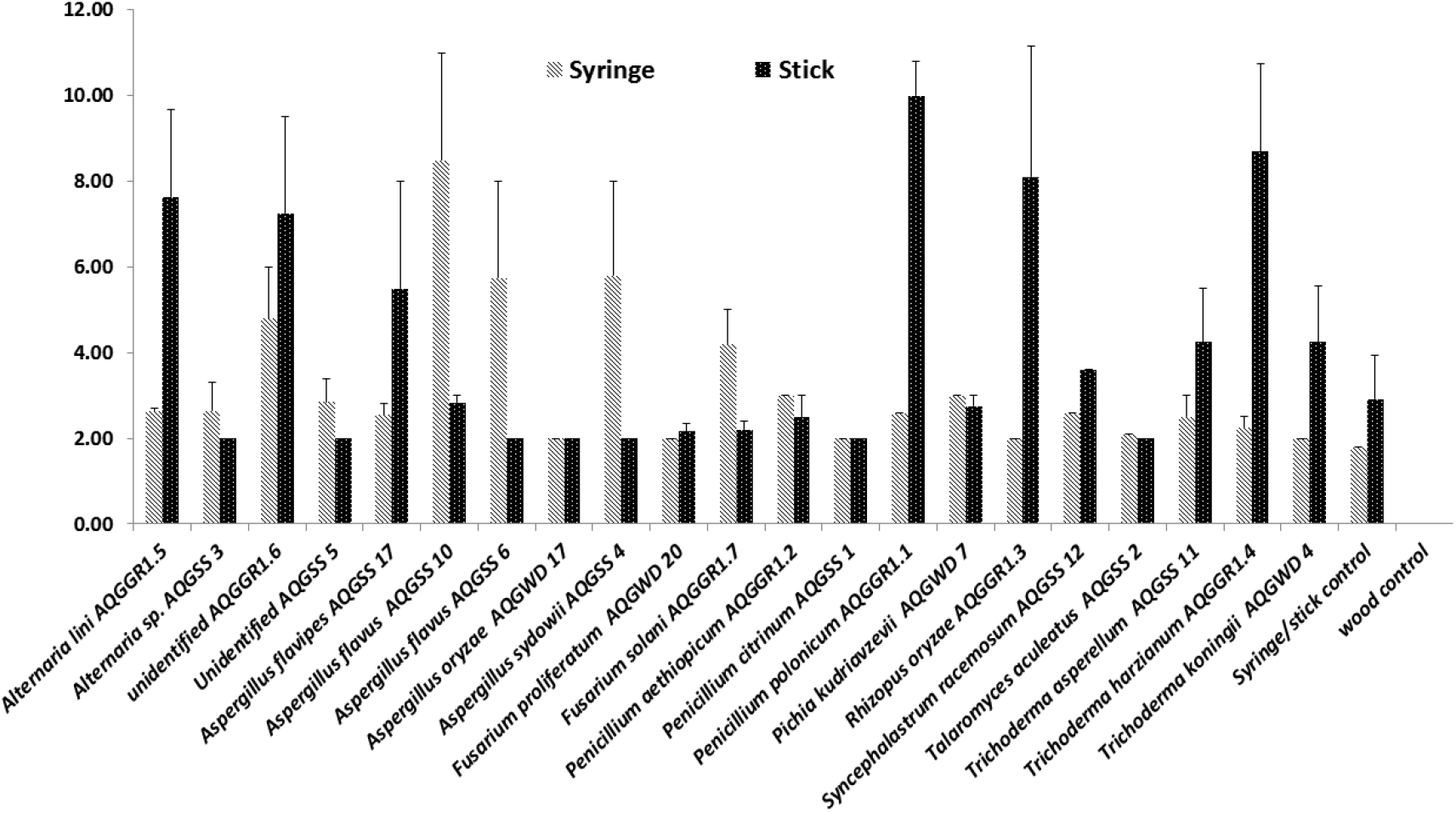
Effect of fungal strain on infection length: The infection length (cm) was measured in two artificial infection methods by different fungal strain after 3 month of inoculation

### 3.2 Effect of fungi on oleoresin production

Twenty one fungal strains were inoculated artificially by stick and syringe method and infected samples were collected after 3 months of inoculation. The oleoresin compounds were extracted by refluxing method. The compounds were identified by Gas chromatography mass spectroscopy is given in Table 1. Agarospirol, Anisyl acetone and Benzyl acetone were identified as major compounds of agarwood oleoresin. It was observed that maximum yield of oleoresin obtained in samples infected by fungal strain *Penicillium polonicum* AQGGR1.1 with 3.2 % in syringe method, while in the case of stick method only 2.05% yield was obtained maximum in *Aquilaria* wood infected by fungal strain *Syncephalastrum racemosum* AQGSS 12. Maximum production of Anisyl acetone was also measured in syringe method in sample infected by *Fusarium solani* AQGGR1.7. Similarly, maximum amount of Benzyl acetone was measured in syringe method, infected by *Penicillium polonicum* AQGGR1.1. In contrast in stick method, 95% organisms showed induction of Benzyl acetone while in syringe method only 66.6% could induce. The marker compound Agarospirol production was measured in 23.8% of samples infected by stick and syringe method. Maximum amount of Agarospirol was measured in *Penicillium polonicum* AQGGR1.1 infected samples. The effect of fungal strains on major compounds induction was analyzed by Venn diagram (Figure 3). In syringe and stick method only 23.8% of fungal strains induced all major compounds, But induction of Benzyl acetone and Anisyl acetone was observed in 71.4% of fungal strains in stick method while only 42.9% in syringe method (Figure 3). Only *Penicillium aethiopicum* AQGGR1.2 found as effective fungus that can induce all major compounds in both method. Infection length was significantly greater in stick method with p value 0.039, while Agarospirol and oil yield was significantly higher in syringe method with p values 0.00 and 0.0024, respectively. There was no significant difference in the two methods in case of Benzyl acetone content.

**Table 1:**
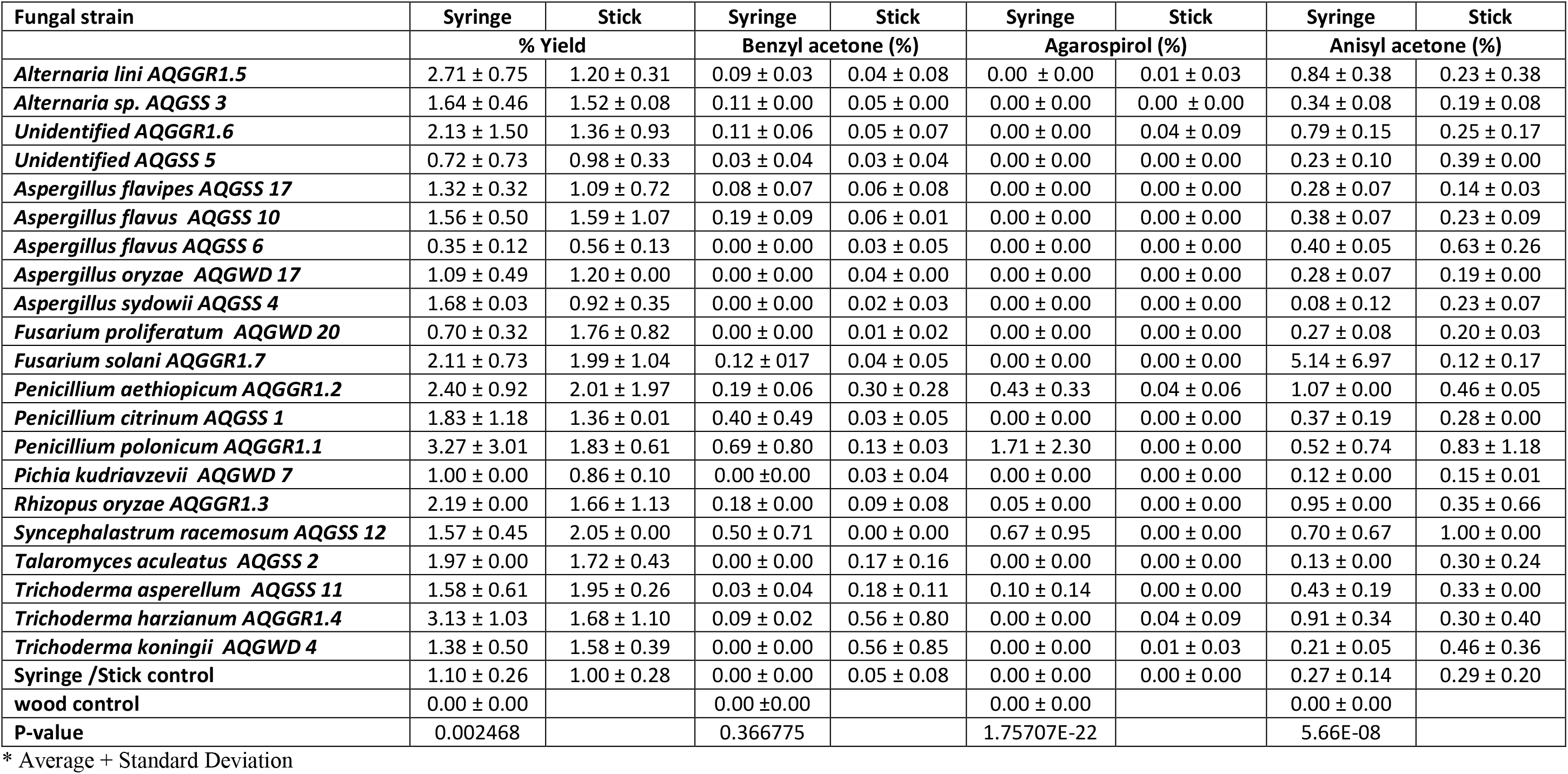
Oleoresin composition of major chemical compounds in samples infected by different fungus

**Figure. 3:**
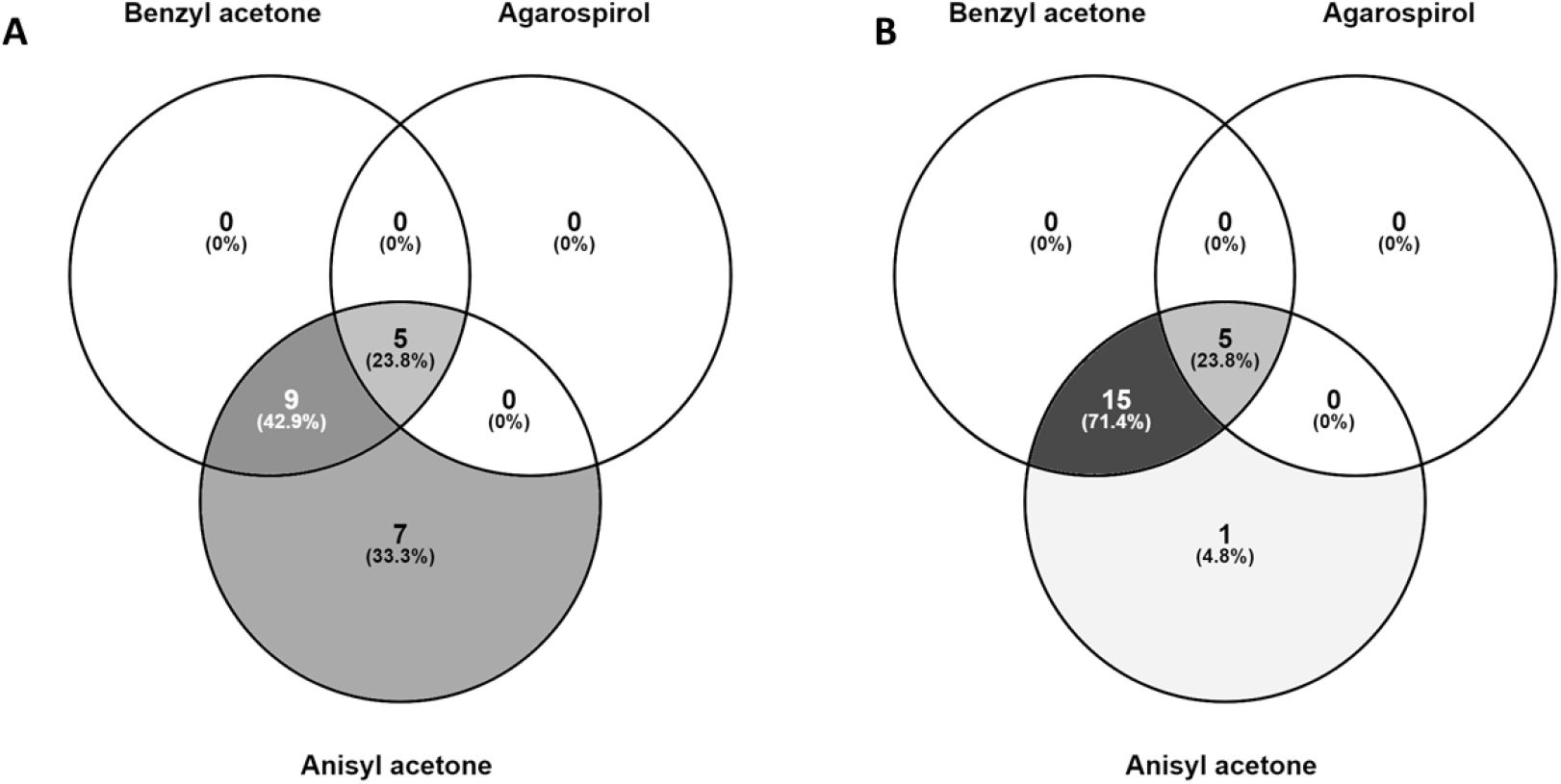
Venn diagram of major compounds induced in (A) Syringe and (B) Stick method

## 4. Discussion

Agarwood is the most expensive wood of the world and its price depends on quality of oleoresin. Oleoresin production is mainly occurred due to defensive action of *Aquilaria* tree against fungal infection. In the current study, we determined the artificial agarwood production potential of twenty one fungal strains belongs to genera *Alternaria, Aspergillus, Fusarium, Penicillium, Pichia, Rhizopus, Syncephalastrum, Talaromyces* and *Trichoderma*. We observed that length of infection is depends on the fungal strain. The infection occurred due to release of cellulolytic and ligninolytic enzyme produced by fungal strains and responsible for disintegration and softening of cellulose, hemicelluloses, pectin, and lignin molecules of plant cell wall (Atri and Sharma 2012). We could find that *Penicillium polonicum* AQGGR1.1 showed maximum infection in stick method up to 10 cm in length. Similarly, Sangareswari et al. (2016) reported that *Aspergillus* isolate AR13 showed maximum cellulase, ligninolytic and laccase activities amongst 17 fungal strains isolated from infected agarwood samples of North Eastern part of India, Tamilnadu and Kerala state of India. The discoloration of wood by lignin degradation has shown as brown or dark wood colour at infected area (Chhipa and Kaushik 2017). Previously, Mohamed et al. (2010) measured only 1.17 cm of infection length after 3 months of artificial inoculation while we could measure up to 10 cm of infection length. The infection was continuously increased more in stick method in comparison to syringe method after next 3 months of agarwood oleoresin production supports the stick method as more infective and efficient artificial infection method. Artificial inoculation of fungi in the *Aquilaria malaccensis* produced the oleoresin after 3 months of infection by both methods. The oleoresin compounds of the infected plant material were analysed by GC-MS and we could find Agarospirol, Anisyl acetone and Benzyl acetone as major chemical compounds. It was hypothesized that if we increase stress to plant it would support more production of oleoresin. The use of bamboo sticks wrapped with fungal mycelia in stick method supposed to produce physical and biological stress to plant for more oleoresin and we observed increased infection length and marker compound Agarospirol in stick method. Stick prevent prolonged from healing to the tree and induce injury continuously and the production of the fragrant resin is associated with wounding and associated fungal invasion (Akther et al. 2013). Sen et al. (2017) reported that *Aquilaria* tree produce the oleoresin as defensive response against fungal attack as pathogen. During fungal plant interaction fungi produce mycotoxins to develop colonization in plant tissue and plant produce chemical compounds for prevention of fungal infection. In our previous study, we observed that presence of Agarospirol is the marker compound showed initiation of oleoresin production due to microbial infection in *Aquilaria malaccensis* (Chhipa and Kaushik 2017). More research is needed to understand the stress development by stick method in continuous production of oleoresin artificially.

## 5. Conclusion

Agarwood oleoresin quality and aroma is dependence on sesquiterpene and chromone type compounds. The amount and chemical composition of oleoresin determined its prices in global market. The production of such compounds can be improved and increased significantly by artificial infection using biological and physical injury. In the present study we have shown combined biological and physical stress in stick method increased infection length and amount of major compounds in comparison to syringe method. *Penicillum* genus could provide potential fungal strains that can be good source of artificial infection in *Aquilaria malaccensis*. Development of stick method as artificial infection method can be helpful in strengthen economy of poor villagers of North Eastern part of India.

## Acknowledgement and/or disclaimers, if any

The work was supported by Department of Biotechnology, Government of India, Research Grant no BT/PR1505/NDB/38/223/2011. We are highly thankful to Mr Surjit Bohra, Golaghat, Assam and R. K. Mohsang Society, Jairampur, Arunachal Pradesh for providing Aquilaria trees for experimental purpose

## Notes

### Competing Interest Statement

The authors have declared no competing interest.

