## Supplementary Table for "Combined effect of biological and physical stress on artificial production of agarwood oleoresin in *Aquilaria malaccensis*"

Table S1: % increase in infection length after next 3 months of development of agarwood oleoresin in both methods

| S.No. | Fungal strain | Syringe method | Stick method |
| --- | --- | --- | --- |
| 1. | ***Alternaria lini AQGGR1.5*** | 0.0 | 0.0 |
| 2. | ***Alternaria sp. AQGSS 3*** | 0.0 | 0.0 |
| 3. | ***Penicillium polonicum AQGGR1.1*** | 0.0 | 5.0 |
| 4. | ***Penicillium aethiopicum AQGGR1.2*** | 0.0 | 0.0 |
| 5. | ***Unidentified AQGGR1.6*** | 0.0 | 98.6 |
| 6. | ***Unidentified AQGSS 5*** | 49.4 | 133.7 |
| 7. | ***Aspergillus flavipes AQGSS 17*** | 0.0 | 0.0 |
| 8. | ***Aspergillus flavus AQGSS 6*** | 0.0 | 0.0 |
| 9. | ***Aspergillus flavus AQGSS 10*** | 0.0 | 47.5 |
| 10. | ***Aspergillus oryzae AQGWD 17*** | 17.8 | 0.0 |
| 11. | ***Aspergillus sydowii AQGSS 4*** | 4.4 | 61.2 |
| 12. | ***Fusarium proliferatum AQGWD 20*** | 17.8 | 0.0 |
| 13. | ***Fusarium solani AQGGR1.7*** | 0.0 | 115.3 |
| 14. | ***Penicillium citrinum AQGSS 1*** | 1.1 | 0.0 |
| 15. | ***Pichia kudriavzevii AQGWD 7*** | 11.1 | 0.0 |
| 16. | ***Rhizopus oryzae AQGGR1.3*** | 17.8 | 0.0 |
| 17. | ***Syncephalastrum racemosum AQGSS 12*** | 17.8 | 106.3 |
| 18. | ***Talaromyces aculeatus AQGSS 2*** | 0.0 | 0.0 |
| 19. | ***Trichoderma asperellum AQGSS 11*** | 0.0 | 0.0 |
| 20. | ***Trichoderma harzianum AQGGR1.4*** | 0.0 | 0.0 |
| 21. | ***Trichoderma koningii AQGWD 4*** | 26.3 | 0.0 |
